# Spermiogenesis alterations in the absence of CTCF revealed by single cell RNA sequencing

**DOI:** 10.1101/2021.03.30.437601

**Authors:** Ulises Torres-Flores, Fernanda Díaz-Espinosa, Tayde López-Santaella, Rosa Rebollar-Vega, Aarón Vázquez-Jiménez, Ian J. Taylor, Rosario Ortiz-Hernández, Olga M. Echeverría, Gerardo H. Vázquez-Nin, María Concepción Gutierrez-Ruiz, Inti Alberto De la Rosa-Velázquez, Osbaldo Resendis-Antonio, Abrahan Hernández-Hernandez

## Abstract

CTCF is an architectonical protein that organizes the genome inside the cell nucleus in almost all eukaryotic cells. There is evidence that CTCF plays a critical role during spermatogenesis as its depletion produces abnormal sperm and infertility. However, the defects produced by the absence of CTCF throughout spermatogenesis have not been characterized. In this work, we performed single cell RNA sequencing in spermatogenic cells without CTCF. We uncovered defects in transcriptional programs that explain the severity of the damage in the produced sperm. At early stages of spermatogenesis, transcriptional alterations are mild. As germ cells go throughout the specialization stage or spermiogenesis, transcriptional profiles become more altered. We found spermatid defects that support the alterations in the transcriptional profiles, and thus we conclude that CTCF depletion alters several transcriptional profiles mostly during spermiogenesis. Our data highlights the importance of CTCF at the different stages of spermatogenesis.

## Introduction

Sperm production in mammals or spermatogenesis involves a complex process of cell division and specialization in order to produce highly specialized germ cells with the only task of fertilizing another germ cell and produce a new organism (White-Cooper and Bausek, 2010). In male gonads, within the seminiferous tubules, the diploid spermatogonia cells undergo mitosis in order to ensure a constant supply of germ line cells. A specific type of spermatogonia commits to the meiotic process, and they duplicated their DNA prior to genetic exchange between homologous chromosomes. After two uninterrupted rounds of meiotic cell divisions, four haploid cells with recombination events are produced (Garg and Martin, 2016). These haploid cells or spermatids enter a differentiation process that will prepare them for their final journey to pursue of a female germ cell. During this process known as spermiogenesis there are changes in cell morphology that will permit the classic torpedo-like shape of the mature sperm (Dadoune, 2003; Torres-Flores and Hernandez-Hernandez, 2020).

Round spermatids undergo an elongation process until they ultimately elongate to form mature sperm. During all the steps of spermatogenesis, chromatin structure and thus transcriptional profiles are very dynamic and specific for every stage (Green et al., 2018; Hermann et al., 2018; Jung et al., 2017; Luo et al., 2020). Recent studies in chromatin organization in meiotic and haploid stages have been shown that the three-dimensional organization of the genome is rather different in these two stages. At meiotic stages, the genome is organized in meiosis-specific TADs (Luo et al., 2020), whereas in haploid cells, the genome is completely reorganized giving rise to a sperm specific epigenome, which is necessary to recapitulate chromatin structure during the embryo development (Jung et al., 2017; van de Werken et al., 2014). It has been shown that proper histone retention in mammal’s sperm has a role in inter-and transgenerational epigenetic inheritance (Siklenka et al., 2015). Nevertheless, there are almost no insight about the interplay among chromatin organization, establishment of sperm epigenome and maintenance of epigenetic information in the sperm. It has been suggested that genome architectural proteins like CTCF and cohesin complexes plus specific epigenetic marks in retained histones shape the sperm epigenome in mice (Jung et al., 2019). Furthermore, CTCF has been involved in the process of histone retention observed in mature sperm(Hernandez-Hernandez et al., 2016). However, the full impact of CTCF during every step of spermiogenesis has been challenging to analyze. Previously, we generated a conditional Knock-out mouse in which *Ctc*f was deleted (*Ctcf*-cKO) at the beginning of the meiotic stage. In this *Ctcf*-cKO mice, meiosis and spermiogenesis were completed, producing low counts of sperm with defects in morphology and histone retention to some extent. Unfortunately, we were not able to identify the phenotypes produced at every stage of spermatogenesis due to the heterogeneity of cell types in the testis (Hernandez-Hernandez et al., 2016).

In this work, we performed single cell RNA sequencing (scRNA-seq) and identified the defects produced in spermatogenesis without CTCF. We found that transcriptional alterations are mild at the early stages of spermatogenesis. However, we observed significant alterations in transcriptional profiles, biological functions, and morphologic defects throughout spermiogenesis. Overall, this paper provides a detailed characterization of the influence of CTCF at several stages of spermiogenesis and assess their functional implications.

## Results

### Single cell RNA seq reveals different cell clusters in spermatogenesis of *Ctcf*-cKO mouse

We previously reported that conditional deletion of *Ctcf* during mice spermatogenesis produces abnormal sperm (Hernandez-Hernandez et al., 2016). However, the stage of spermatogenesis at which these abnormalities occur have not been elucidated. First, we sought to find out whether spermatogenesis in the testes from *Ctcf*-cKO mouse was fully represented. We isolated and sequenced the RNA of single cells from mouse testes of four mice from which 909 cells passed quality control (QC) (406 cells from wild-type and 503 cells from *Ctcf*-cKO mice). We then computed the Uniform Manifold Approximation and Projection for Dimension Reduction (UMAP) (Figure 1A) to visualize data distribution and identify cell subpopulations. As a result of this analysis, we noticed that some cell populations from both conditions clustered in the same phenotypic space; while other populations specific to the *Ctcf*-cKO mouse, clustered independently within the UMAP embedded space (Figure 1A). To evaluate the existence and identity of cell subpopulations we pooled the data and performed unsupervised clustering with Seurat package. Overall, we identified 11 clusters throughout both conditions (Figure 1B). We observed that clusters 3 and 5 are specific for *Ctcf*-cKO spermatogenesis whereas cluster 4 is absent in these mice (Figure 1B).

**Figure 1.**
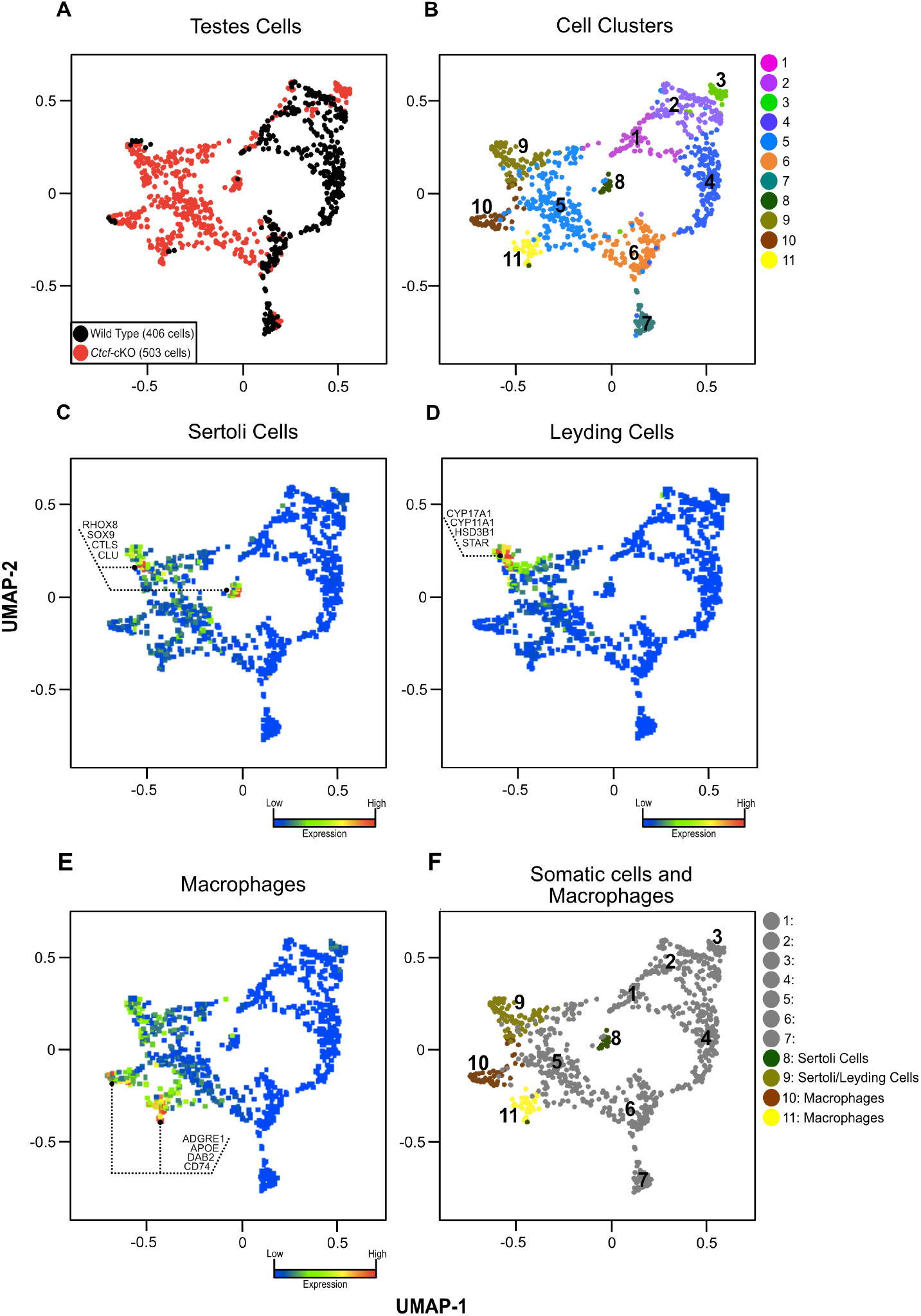
Dimensionality reduction and clusters identification. (**A**) UMAP visualization of testes cells from wild type (black dots) and *Ctcf*-cKO (red dots) mice. (**B**) Clustering of cell populations across both conditions. A different color has been assigned to each cluster. (**C**) Molecular markers used to identify Sertoli cells, (**D**) Leyding cells and (**E**) Macrophages. The scale color represents the sum of the expression levels per cell of the listed genes across all UMAP visualization. (**F**) Overlay of clusters with somatic cells and macrophages.

Since conditional depletion of *Ctcf* takes place in cells that undergo spermatogenesis (Hernandez-Hernandez et al., 2016), we searched for cells that will not enter this process and excluded them from further analysis. By identifying the expression (Log 2-Fold change >1.5, p value<0.05) of molecular markers that have been previously identified by scRNA-seq (Green et al., 2018; Hermann et al., 2018; Suzuki et al., 2019), we classified clusters 8-11 as Sertoli cells, Leyding cells and macrophages respectively (Figures 1C-F). Therefore, the remaining clusters 1-7 might represent spermatogenesis in both conditions. However, the presence of the *Ctcf*-cKO specific clusters 3 and 5, and the absence of cluster 4, suggest that *Ctcf* deletion induce major changes in the transcriptome of these spermatogenic clusters. To corroborate that *Ctcf*-cKO mice did not express the mRNA of CTCF during spermatogenesis, we looked for its expression profile in our UMAP and only observed its expression in WT spermatogenic cells (see also Figures S1, A and B).

### Cells clusters recapitulated spermatogenesis

To identify cells in the main stages of spermatogenesis (i.e., mitosis, meiosis and spermiogenesis), we mapped the expression of stage-specific markers based on previous reports (Green et al., 2018; Hermann et al., 2018). We found mitotic cell markers preferentially expressed in cluster 1 (Figures 2A, 2B and see also Figure S1C), while meiotic cell markers were found in clusters 1 and 2, (Figures 2A and 2C). These two clusters are present in both WT and *Ctcf*-cKO mice.

**Figure 2.**
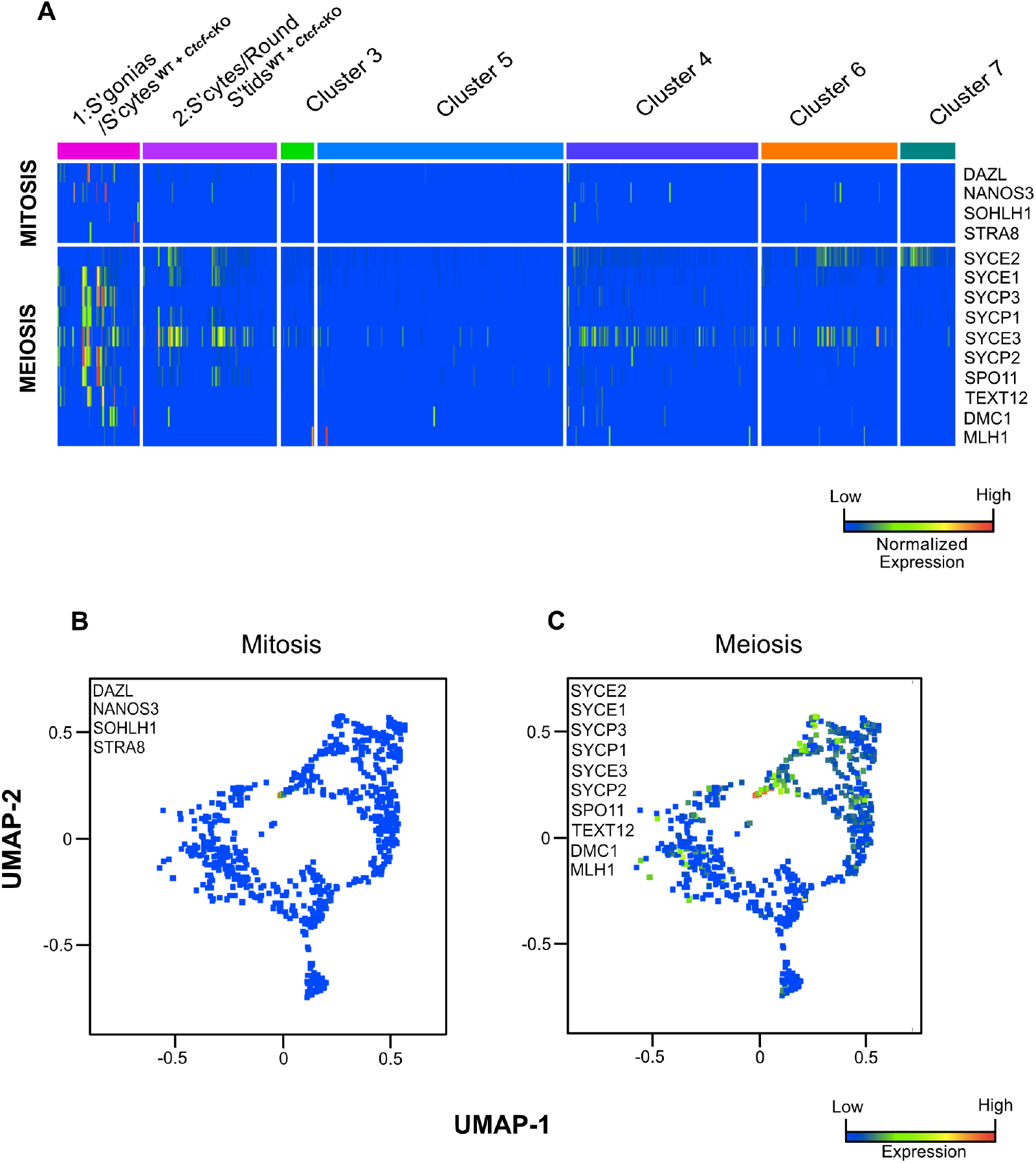
Identification of clusters with mitotic and meiotic cells. (**A**) Heat map displaying the expression levels of the markers used to identify mitotic and meiotic cells. (**B**) UMAP projection with the expression of the mitotic molecular markers in the cell clusters. (**C**) UMAP with the expression of the meiosis molecular markers in the cell clusters. In (**A**), the scale color displays the <u>mode normalized expression</u> of each gene identified in each cluster. <u>Mode Normalizatio</u>n means maximum values are shown in red, minimum in blue, all values reported are relative, per gene, regardless of absolute transcript count, while in figures (**B** and**C**) the scale color shows the total sum expression count of genes within the listed gene-set, per cell across UMAP visualization.

We next searched for clusters with spermiogenesis markers and observed that clusters 3 to 7 express them (Figures 3A and 3B). It has been reported that only round spermatids display gene transcriptional levels that are detectable by scRNA-seq studies. Based on this detectable transcription, spermatids have been divided into early-, mid- and late-round spermatids (Green et al., 2018; Hermann et al., 2018). To identify these stages of round spermatids we looked into the normalized expression of their markers (early: ACRV1, TSSK1, GM1035, SPEER4E; mid and late: PRM1, PRM2, TNP1, TNP2, HSPA1L, ACTL7B, LRRC34, SPACA1, CYP2A12, CD63 and 4930513O06RIK) (Fig. 3A). Expression profile of genes involved in histone replacement (i.e., PRM1, PRM2, TNP1 and TNP2) (Torres-Flores and Hernandez-Hernandez, 2020), revealed that cluster 4, 6, and 7 correspond to early-, mid- and late-round spermatids, respectively (Figure 3A). Furthermore, we overlayed the global expression of these spermiogenesis markers in our UMAP projection and found that cluster 6 (mid-round spermatids) has the highest sum expression count of these genes in comparation to other clusters (Figures 3B and 3C).

**Figure 3.**
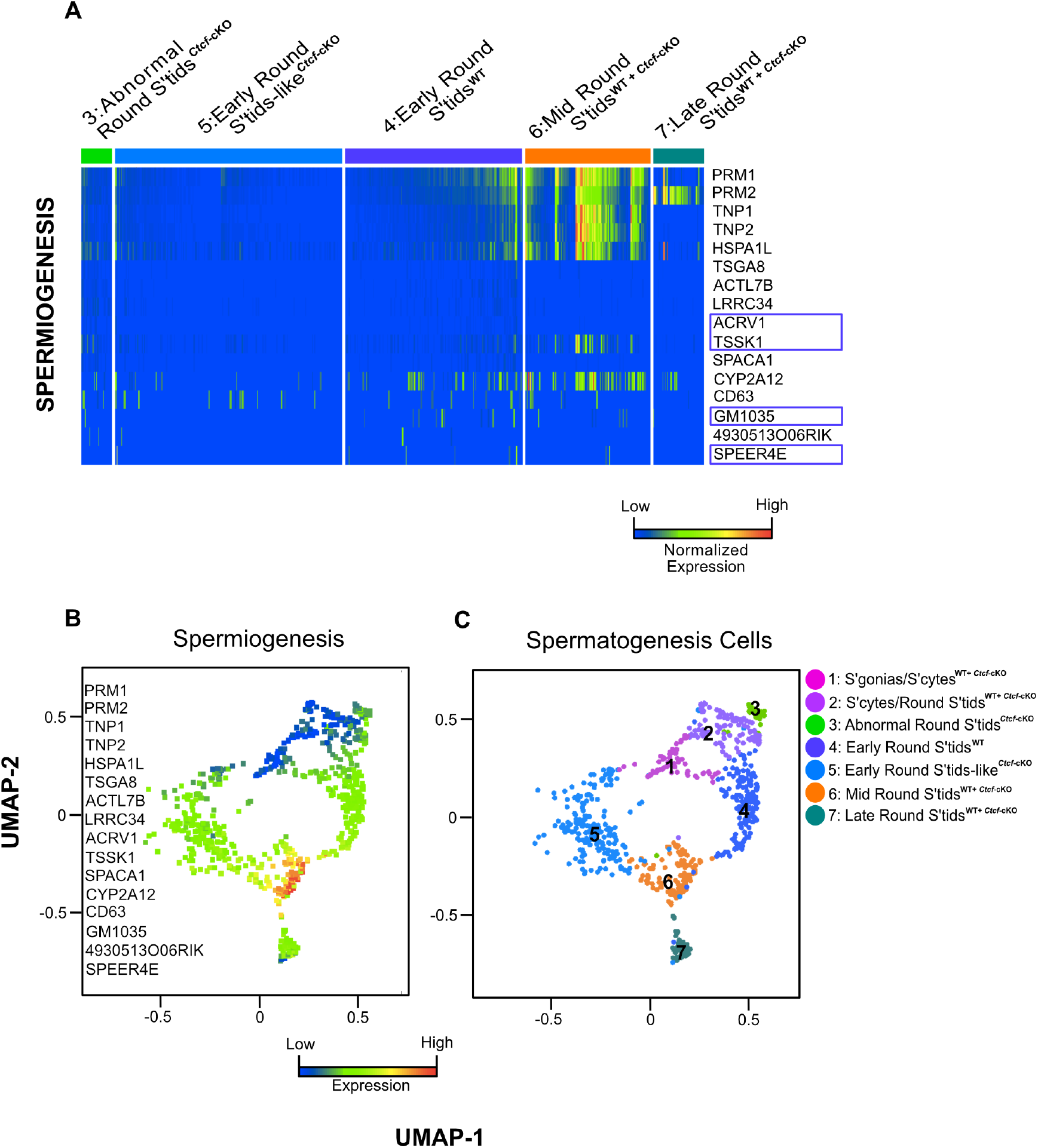
Identification of clusters with cells in spermiogenesis. (**A**) Heat map displaying the expression levels of the markers used to identify cells in spermiogenesis. Dark blue rectangle highlights early round spermatids genes, while genes without rectangle correspond to mid and late round spermatids. (**B**) UMAP projection with the expression of the spermiogenesis molecular markers in the cell clusters. (**C**) UMAP projection with the expression of the spermatogenesis molecular markers in the cell clusters. In (**A**), the scale color displays the <u>mode normalized expression</u> of each gene identified in each cluster. Mode Normalization means maximum values are shown in red, minimum in blue, all values reported are relative, per gene, regardless of absolute transcript count, while in (**B** and **C**) the scale color shows the total sum expression count of genes within the listed gene-set, per cell across UMAP visualization.

Although, cluster 4 is only found in WT, clusters 6 and 7 contain cells from WT and *Ctcf*-cKO mice. These data indicate that at least for WT, spermiogenesis progresses from cluster 4, towards cluster 6 and ends at cluster 7. In contrast, the *Ctcf*-cKO spermiogenesis, progression seems parallel to WT but without early round spermatids. Therefore, we sought to find early round spermatids-like cluster(s) in the *Ctcf*-cKO specific clusters. Despite mapping of round spermatids markers to cluster 3 and 5 (specific for *Ctcf*-cKO) their identification into early, mid and late spermatid populations is not clear (Figure 3A). Hence, we assessed the normalized gene expression of round spermatids markers in all the clusters. We observed that gene expression distribution of the early round spermatid markers (TSGA8, ACTL7B, TSSK1 and LRRC34) and mid- to late-round spermatid markers (TNP1, TNP2, PRM1 and PRM2) in cluster 5 are more similar to cluster 4 (early round spermatids) and significantly different from those expression distributions seen in cluster 6 (mid round spermatids) (see also Figure S2). Taking into account the global expression profiles of spermiogenesis markers (Figure 3B), and the normalized gene expression distribution (see also Figure S2) we identified cluster 5 as early-round spermatids-like specific for *Ctcf*-cKO mouse. Furthermore, despite displaying expression of spermatids markers (Figures 3A and 3B), the expression distribution reveals that cluster 3 represents a group of spermatids with altered expression of these markers (see also Figure S2). Thus, we recognized this cluster as abnormal round spermatids specific for *Ctcf*-cKO mouse. Altogether, our analyzes helped us to identify each one of the clusters present in the UMAP representations from both *Ctcf*-cKO and WT spermatogenesis (Figure 3C). Furthermore, our data indicate that transcriptional profiles of spermiogenesis in the *Ctcf*-cKO are drastically altered at the early-round spermatid stage at to a lesser extent in mid and late round spermatids.

### Pseudotime analysis reveals a different trajectory but similar outcome in *Ctcf*-cKO spermiogenesis

With the Identification of spermatogenesis clusters in WT and *Ctcf*-cKO spermatogenesis we suggest that progression moves from cluster 1 to 2 (from spermatogonia to meiotic cells) in both conditions. Early round spermatids in WT are in cluster 4 whereas early round spermatids-like are in cluster 5 in *Ctcf*-cKO mice. Finally, mid- and late round spermatids are in clusters 6 and 7 respectively in both conditions. In order to identify a possible progression based on their profiles of gene expression of mitotic, meiotic, and spermiogenesis specific genes (Figure 4A), we performed Monocle Dimensionality Reduction (MDR) and pseudotime analysis using Monocle 2 (Trapnell et al., 2014). We detected five stages of progression in both, the WT and *Ctcf*-cKO spermatogenesis (Figures 4B and 4C). Since we did not observe clear differences between the WT and *Ctcf-*cKO, we overlaid these stages into our UMAP projection and noticed that the stages of progression are analogous to the biological progression observed in these UMAPs (i.e., mitosis towards meiosis and meiosis towards spermiogenesis) in both conditions (Figures 4D and 4E).

**Figure 4.**
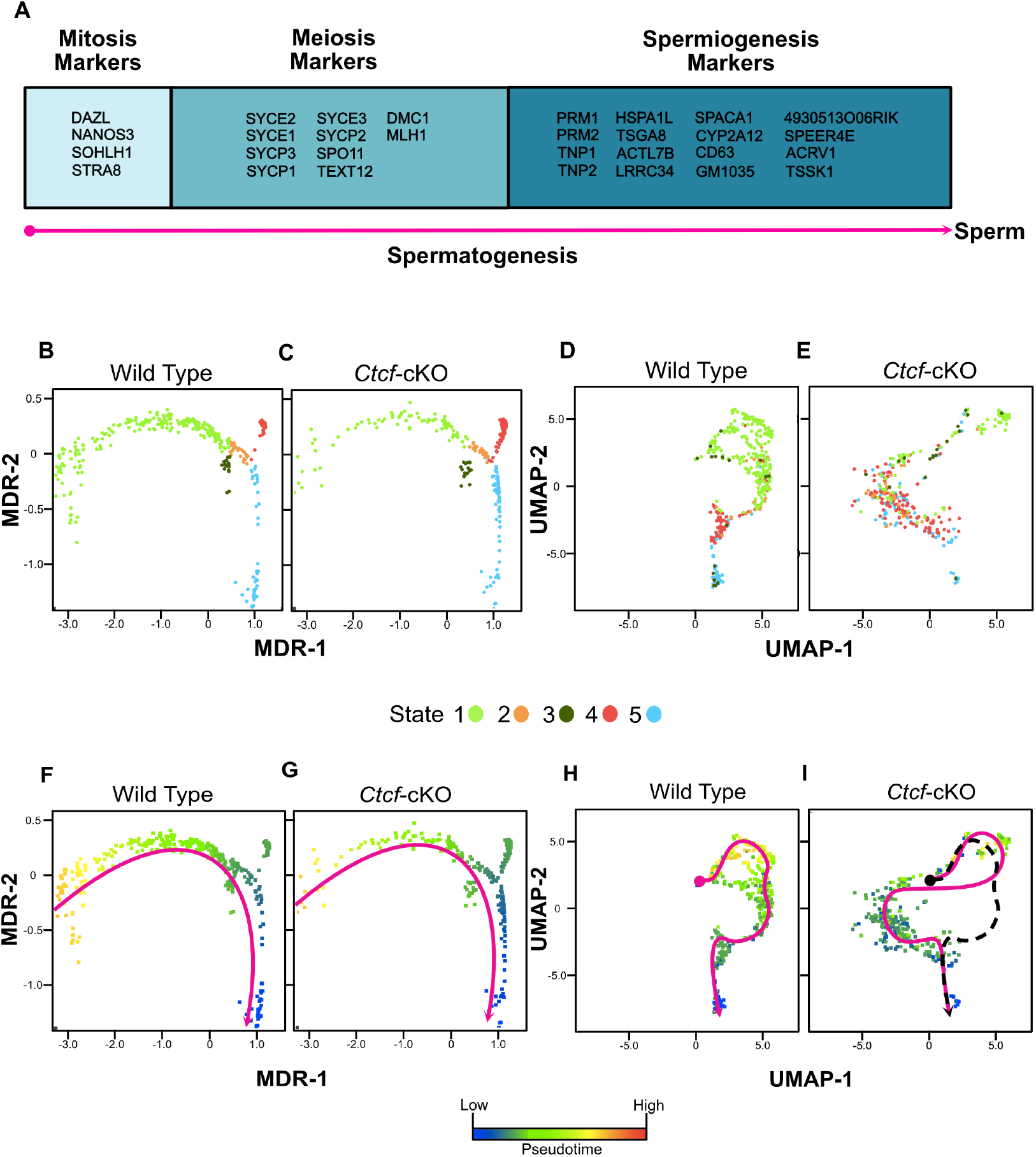
Spermatogenesis Pseudotime Analysis. (**A**) Genes expressed across the three major steps of spermatogesis: mitosis, meiosis and spermiogenesis. (**B**) and (**C**) Monocle dimentionally reduction states (MDR) identified in Wild type and *Ctcf*-cKO cell clusters. Each dot represent a cell while each color represent an identified state. (**D**) and (**E**) Overlapping of MDR states into UMAP projections of Wild type and *Ctcf*-cKO datasets. (**F**) and (**G**) Pseudotime progression of cells in spermatogenesis. Key color scale indicates the development prediction from blue to red, representing low to high expression respectively. Magenta curved arrow designates the prediction trajectory in our model. (**H**) and (**I**) Overlapping of MDR pseudotime into UMAP projections of Wild Type and *Ctcf*-cKO datasets. Magenta curved arrows point out the trajectory prediction of spermatogenesis in Wild type and *Ctcf-*cKO, respectively. In (**I**), the dotted curved arrow represents the predicted trajectory in Wild Type spermatogenesis. Overlapping of both curves shows the same beginning and end of the spermatogenesis in both genetic backgrounds but a different trajectory in *Ctcf*-cKO spermatogenesis.

We then overlaid the pseudotime projection on our MDR plot and found that in the WT the highest transcription rates of our markers matched with the spermatogonias/spermatocytes clusters. Furthermore, transcription rates dropped gradually following the dynamics of spermatogenesis until they ceased at the late round spermatids cluster (Figures 4F and 4G). Although early round spermatids are in a different and unique cluster in the *Ctcf-*cKO, our pseudotime analysis predicted the same trajectory from spermatogonia towards spermatocytes prior early, mid and late round spermatids (Figures 4H and 4I), suggesting that spermiogenesis was a continuous process in the absence of CTCF, despite morphological and biochemical alteration during the whole process.

### Transcriptional profiles are drastically altered in early round spermatids of *Ctcf*-cKO spermatogenesis

Our previous analyses suggest that early-round spermatids experience the greatest damage in the absence of CTCF, but also that they progress towards mid- and late-round spermatids. To assess the alteration in transcriptional signatures across spermatogenesis we performed gene expression analysis between *Ctcf*-cKO and WT cells clusters (fold change greater than 2, p-value and FDR: 0.05; see also Figure S3). Then, with the DataBase for annotation visualization and integrated discovery 6.8 version (DAVID 6.8), we analyzed the up-and down-regulated genes. The entire list can be found in Table S1 and Table S2. We selected the top-five GO terms results and we found that across all the clusters the down-regulated genes in the *Ctcf*-cKO spermatogenesis have annotations related to proper progression of spermatogenesis, sperm development and male fertility (see also Figure S4). On the contrary, the up-regulated gene annotations are related to cell process not related to spermatogenesis (see also Figure S4).

Furthermore, we sought to perform a more robust characterization of the overrepresented altered biological pathways throughout the spermatogenesis in the *Ctcf*-cKO mouse, thus we performed pathway enrichment analysis using the GSEA tool (Subramanian et al., 2005) and used the Cytoscape software for visualization of enriched pathways (represented as nodes of sizes that are proportional to the number of genes mis-regulated in the specific gene set). When several pathways are associated by similarity they are displayed as interconnected within circles, representing higher-level processes given a specific biological function. In the cluster of spermatogonia and spermatocytes (cluster 1), we found some isolated nodes related to genes specifically expressed in male sexual organs and differentially expressed genes in juvenile *Spo11*-KO testes (Figures 5A-C; see also Table S3), supporting our finding that this cluster contains specifically spermatogonia and spermatocytes; since in *Spo11*-KO mice, spermatogenic cells do not progress beyond early spermatocytes (Smirnova et al., 2006). In the cluster of spermatocytes and round spermatids (cluster 2), we also observed a few isolated nodes related to genes specifically expressed in male sexual organs and down-regulated genes in *Cdx* mutant mice that are important for embryonic development (van Nes et al., 2006) (Figure 5C, see also Table S4).

**Figure 5.**
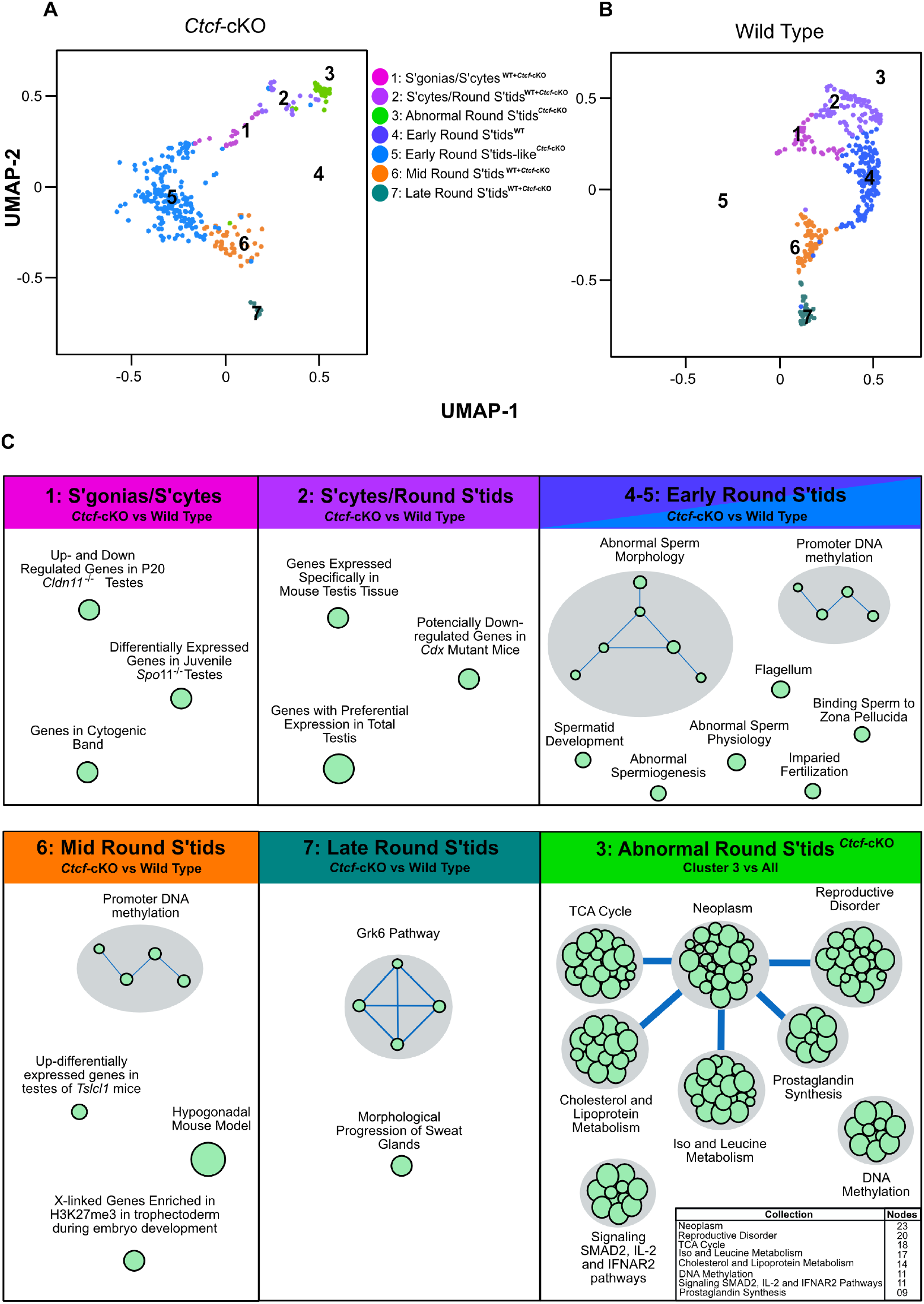
Gene Set Enrichment Analysis (GSEA) of *Ctcf*-cKO vs Wild type spermatogenesis. (**A**) and (**B**) showed clusters and their identity in UMAP visualizations of spermatogenesis cells from Wild type and *Ctcf*-cKO mice, respectively. (**C**) Positively enriched maps of biological functions altered in the different stages of spermatogenesis from *Ctcf*-cKO mice. Each green circle (node) represents a pathway, circle size represents the number of genes with altered transcriptional activity in each pathway. Link between pathways was established when 2 pathways had a Jaccard index > 0.4, given the expressed genes. The embedded table in last inset in (**C**), shows the number of pathways enriched in each collection. All the enriched pathways have an FDR < 0.05 and a p value < 0.01.

In the cluster of early round spermatids (clusters 4 and 5 in WT and *Ctcf*-cKO, respectively), we observed isolated nodes but also two biological functions represented by several nodes (Figure 5C). The isolated nodes are associated with abnormal spermatogenesis, spermatid differentiation, sperm physiology, and impaired fertilization. Whereas the biological functions with several nodes are associated with abnormal sperm morphology and promoter DNA methylation (Figure 5C; see also Table S5). In the cluster of middle round spermatids (cluster 6), we also found the biological function of promoter DNA methylation to be overrepresented. Meanwhile, isolated nodes are associated with genes expressed specifically in mouse testis, hypogonadal mouse model (apoptosis increase) (Chausiaux et al., 2008), and up-regulated genes in testes of *Tslc1*-KO mice (Yamada et al., 2006) (Figure 5C; see also Table S6). Then, in the cluster of late round spermatids (cluster 7), we found the biological function of the Grk6 pathway, that has been reported to be regulating engulfment of apoptotic cells (Nakaya et al., 2013) (Figure 5C; see also Table S7).

Finally, in cluster 3 of abnormal round spermatids (that is unique of the *Ctcf*-cKO mice), we found several biological functions enriched (Figure 5C; see also Table S8). Not all these biological functions are directly involved in hallmark spermatogenesis processes as they comprise diverse biological functions like TCA cycle, Neoplasm, Reproductive disorder, Cholesterol and lipoprotein metabolism, Iso- and leucine metabolism, DNA methylation, prostaglandin synthesis and Signaling SMAD2, IL-2 and IFNAR2 pathways (Figure 5C; see also Table S8). Altogether, our characterization of the overrepresented altered biological pathways throughout the spermatogenesis in the *Ctcf*-cKO mouse, reveals that early round spermatids suffer the greatest damage as they display the largest number of biological pathways altered in the *Ctcf*-cKO mice.

### Spermatids in *Ctcf*-cKO mice display abnormalities in acrosome formation and nuclear compaction

Previously, we reported that elongated spermatids and mature sperm display chromatin compaction defects in the cell nuclei (Hernandez-Hernandez et al., 2016). However, our present analyzes identified a drastic transcriptome alteration at the stage of early round spermatids and onwards in the *Ctcf*-cKO spermatogenesis. Thus, we decided to perform a detailed morphological characterization of cells in spermiogenesis at Electron Microscopy (EM) level. We found mainly two phenotypes in spermatids that support the abnormal sperm morphology reported in the GSEA analysis.

The first one is related to acrosome biogenesis. In round spermatids from WT mice, we observed the acrosome sac and the head cap (Hc) with their expected spherical formation on the surface of the nucleus (Clermont, 1993), whereas in the *Ctcf*-cKO round spermatids these two structures are rather flattened (Figures 6A and 6B). Following spermatids differentiation from spherical to an asymmetric shape in steps 8-10, we observed that the acrosome and the Hc are condensed and flattened, giving rise to a highly electron-dense acrosome vesicle (Kierszenbaum et al., 2003; Meistrich, 1993). However, we did not observe these changes in the formation of the acrosome vesicle in the *Ctcf*-cKO elongating spermatids (Figures 6C and 6D). Furthermore, in elongated spermatids from WT mice, we observed a well-defined acrosome vesicle in the apical region of the elongated nucleus, whereas in the elongated spermatids from *Ctcf*-cKO we observed a non-well-defined acrosome vesicle in other regions of the nucleus rather than on its apical part (Figures 6E and 6F). Finally, we observed that in the mature sperm from the cauda epididymis of *Ctcf-*cKO mice the acrosome did not form properly (Figures 6G and 6H).

**Figure 6.**
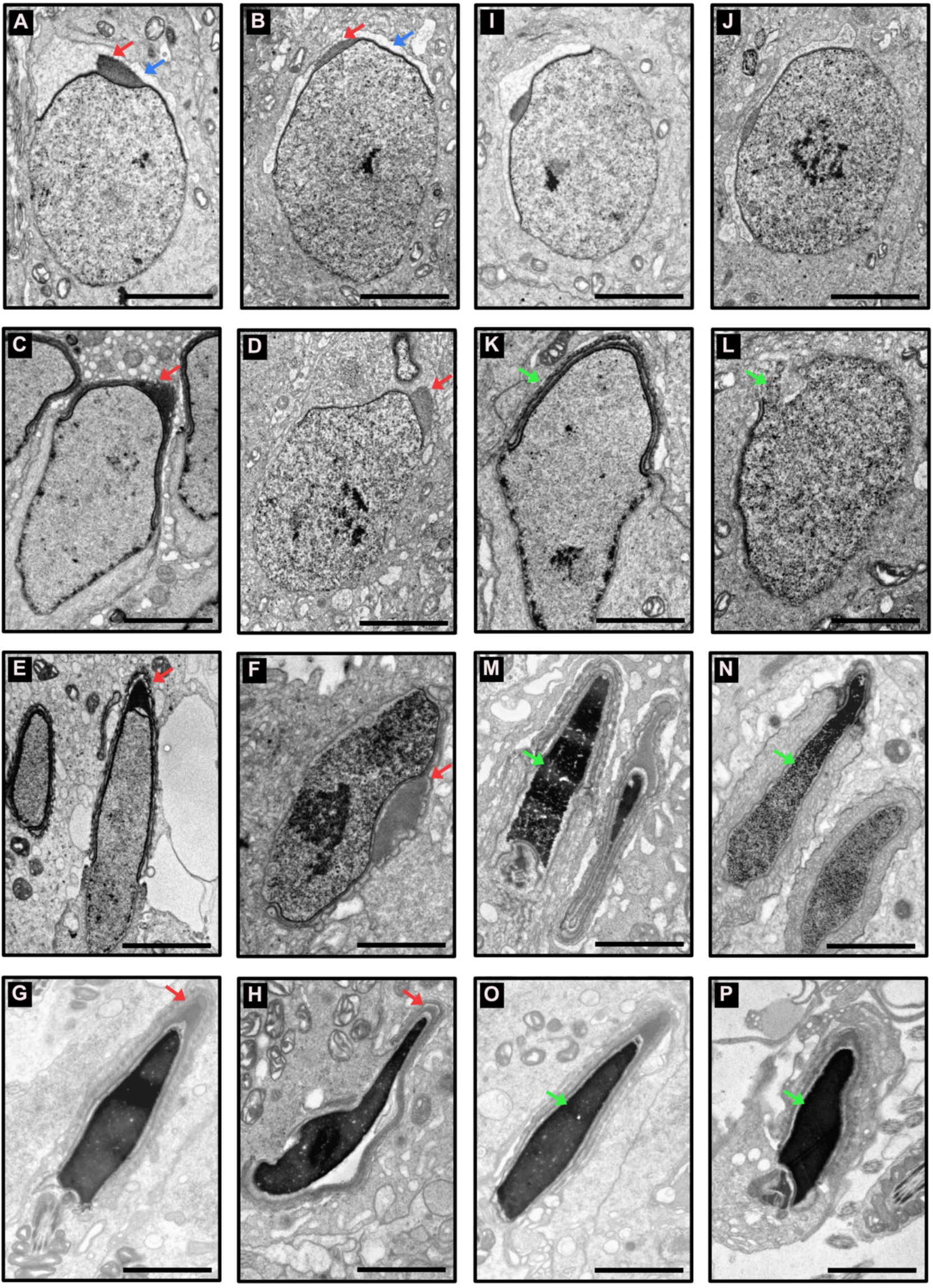
Electron microscopy of spermatids from Wild Type and *Ctcf*-cKO mice. (**A**) shows acrosome sac (red arrow) and head cap (blue arrow) from a WT spermatid. (**B**) Abnormal acrosome sac (red arrow) and head cap (blue arrow) from a *Ctcf*-cKO spermatid. (**C**) Flattened and condensed head cap (red arrow) from a WT elongating spermatid. (**D**) Abnormal head cap (red arrow) from a *Ctcf*-cKO elongating spermatid. (**E**) Properly formed acrosome in apical anterior end of the elongated spermatids from WT mice. (**F**) Mispositioned acrosome (red arrow) in an elongated spermatid from *Ctcf*-cKO mice. (**G**) Acrosome in the apical anterior end of a mature sperm from WT mice. (**H**) Abnormal acrosome and mature sperm morphology from *Ctcf*-cKO mice. (**I**) and (**J**) Proper nuclear morphology and chromatin compaction in round spermatids from WT and *Ctcf*-cKO mice, respectively. (**K**) Elongating spermatid with electron-dense material associated to the nuclear envelope (green arrow) from WT mice. (**L**) Elongating spermatid with discontinuities in the dense structure associate to the nuclear envelope and in the nuclear envelope itself (green arrow) from *Ctcf*-cKO mice. (**M**) Elongated spermatid with condensed chromatin (green arrow) from WT mice. (**N**) Elongated spermatid with abnormal chromatin condensation from *Ctcf*-cKO mice. o Mature sperm displaying full chromatin packing inside the nucleus (green arrow) from WT mice. p Mature sperm displaying apparently full chromatin packing (green arrow) but with abnormal head morphology.

The second phenotype that we underscored was in nuclear shape. We observed nuclear shape defects in the spermatids of *Ctcf-*cKO mice starting at the stage of elongating spermatids. At this stage, we observed defects in the continuity of the nuclear envelope and nuclear shape (Figures 6K and 6L). Meanwhile, in elongated spermatids we observed abnormalities in chromatin compaction (Figures 6M and 6N), we observed abnormal nuclear shape in mature sperm (Figures 6O and 6P).

### The protamine incorporation pathway is affected in *Ctcf-*cKO spermatids

In mature sperm of *Ctcf*-cKO mice, PRM1 is detected in lower amount compared to WT mice. However, alteration at the transcriptional levels during the different stages of spermiogenesis have not been reported (Hernandez-Hernandez et al., 2016). Since cluster 6 has the highest transcriptional levels of most of the protamine incorporation players (i.e., TNP1, TNP2, and PRM1) (Figures, 3A and 7A), we searched for transcriptional changes in these factors and observed lower expression levels of TNP1 and TNP2 in the *Ctcf*-cKO mice (Figures 7A and 7B). However, by immunofluorescent assays, we observed that whereas in elongated/elongating spermatids on WT mice, both TNP1 and TNP2 proteins are co-expressed (Figures 8A-D), while, in 25% of the analyzed seminiferous tubules from *Ctcf*-cKO mice, some spermatids without the classical elongated shape display only TNP1, but not TNP2 (Figures 8E-H). Furthermore, elongated spermatids from both WT and *Ctcf*-cKO display PRM2 and PRM1 (Figures 8I-P) and (Figures 9A-I), respectively. However we found that 30% of the analyzed seminiferous tubules from *Ctcf-*cKO mice display “spermatids” with round-like shape that express PRM1 (Figures 9 G-H). These data show that the protamine incorporation process is altered in *Ctcf*-cKO spermatids. Additionally, it sems that in a sub population of spermatids in the *Ctcf*-cKO either cell morphology or transcriptional programs are completely different to the typical elongating/elongated spermatids. In any case, we found and corroborated that the protamine incorporation mechanism is dysregulated in mid-round spermatids (at the transcriptional level) and in elongating/elongated spermatids (at the protein level) in the spermiogenesis of *Ctcf-*cKO mice.

**Figure 7.**
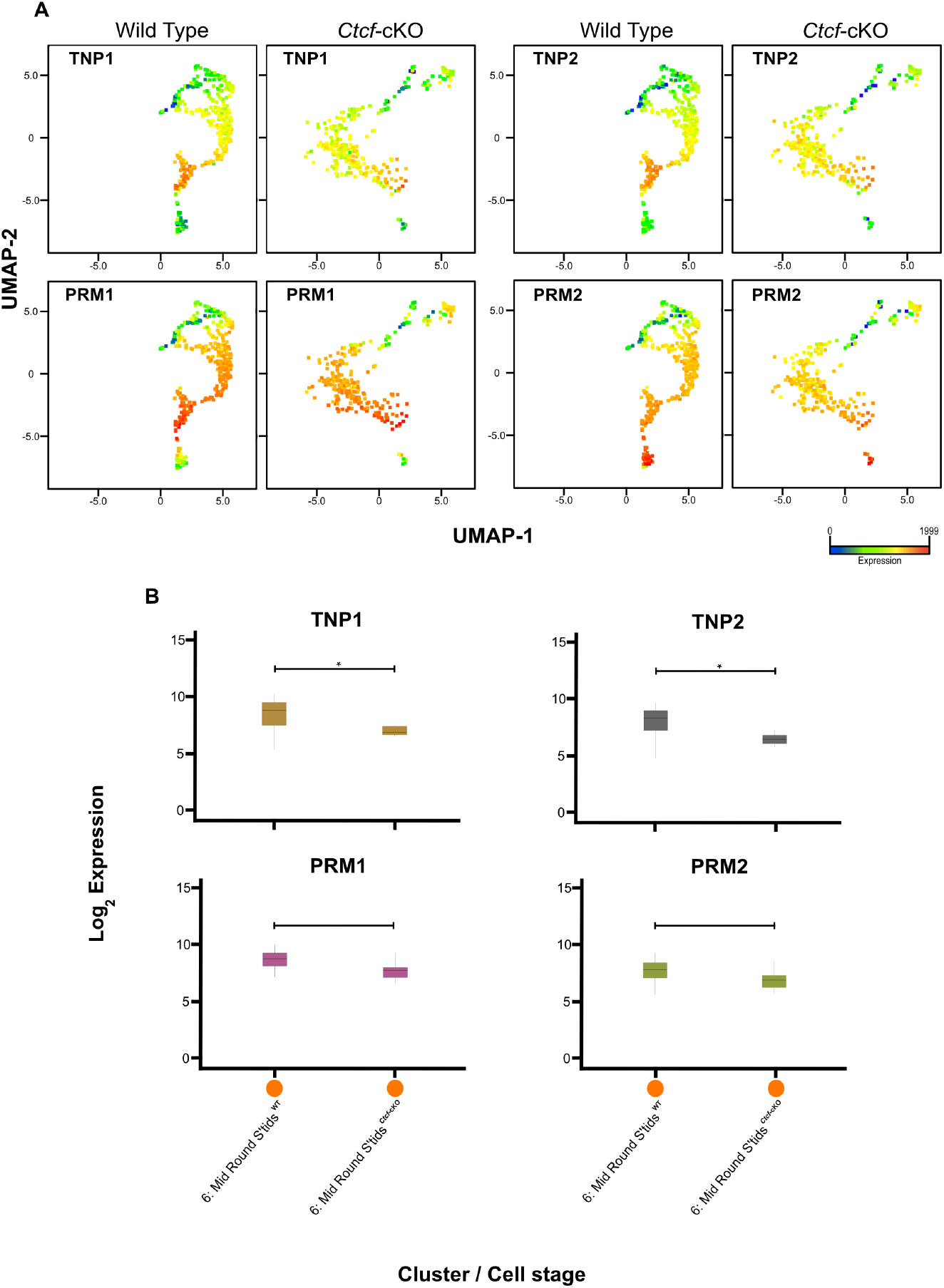
Transcriptional expression profiles in mid-round spermatids from WT and *Ctcf*-cKO mice. (**A**) UMAP projection displaying the expression patterns of TNP1, TNP2, PRM1 and PRM2 in the cluster of mid-round spermatids (cluster 6) from WT and *Ctcf*-cKO mice. (**B**) Expression levels of TNP1, TNP2, PRM1 and PRM2 in mid-round spermatids from WT and *Ctcf*-cKO mice. TNP1 and TNP2 display significantly less expression levels in the *Ctcf-* cKO spermatids (Kruskal-Wallis test, p<0.05). The scale color in (**A**), represents the expression level of each transcript per cell.

**Figure 8.**
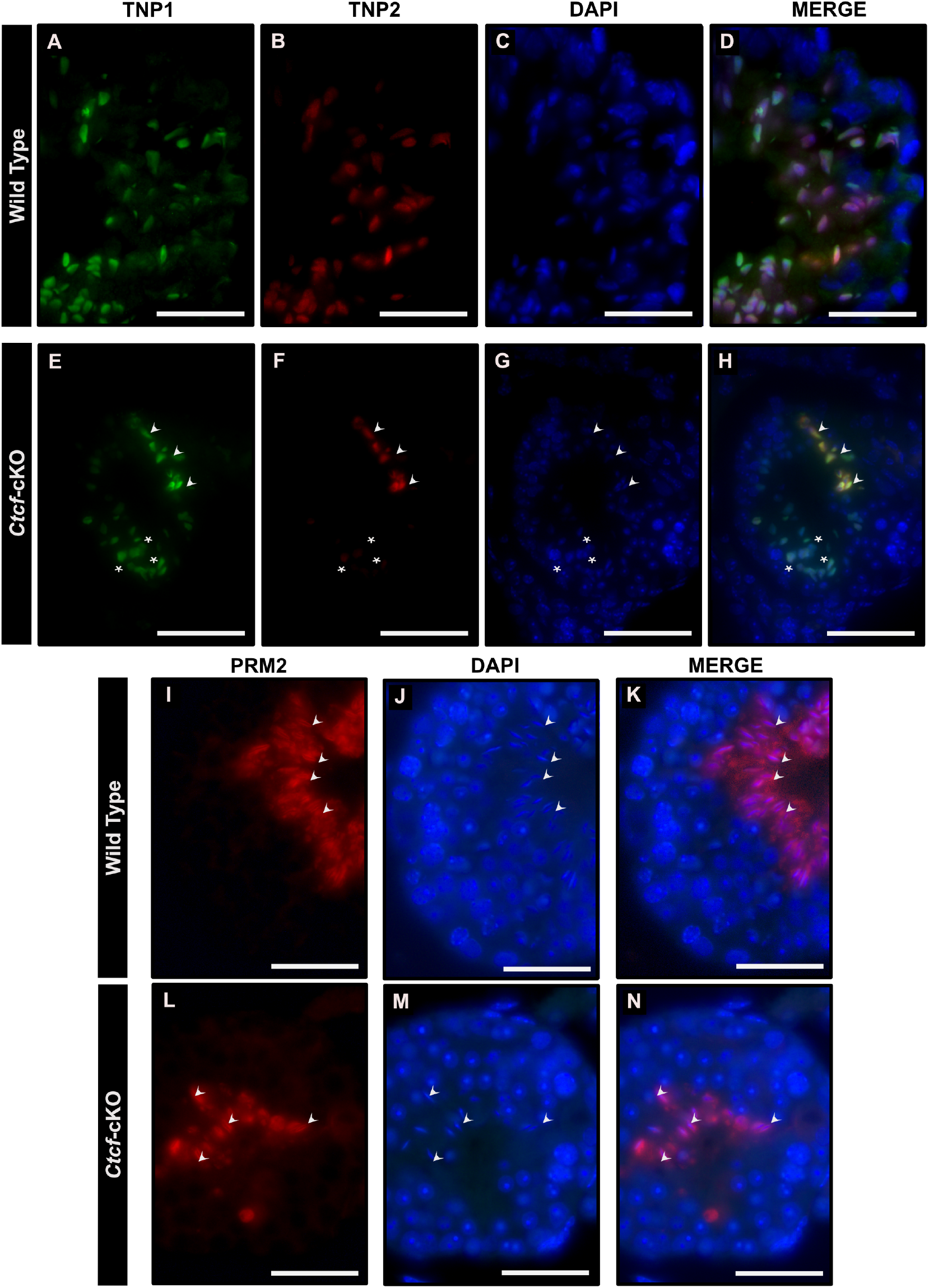
Immunofluorescent staining pattern of TNP1, TNP2 and PRM2 in Wild Type and *Ctcf*-cKO testis sections. (**A-D**) Show simultaneous immunodetection of TNP1 and TNP2 in elongated spermatids from WT testis sections. (**E-H**) Simultaneous immunodetection of TNP1 and TNP2 in elongated spermatids from *Ctcf*-cKO testis sections. Elongated spermatids with immunodetection of both proteins are displayed with arrows, whereas abnormal spermatids with signal only for TNP1 are displayed with asterisks. 25% of the analyzed seminiferous tubules displayed these abnormal spermatids with abnormal staining pattern. Three mice of each genotype were analyzed. (**I-K**) Immunodetection of PRM2 in elongated spermatids from WT testis sections. (**L-N** Immunodetection of PRM2 in elongated spermatids from *Ctcf*-cKO testis sections. Scale bars represent 25 micrometers.

**Figure 9.**
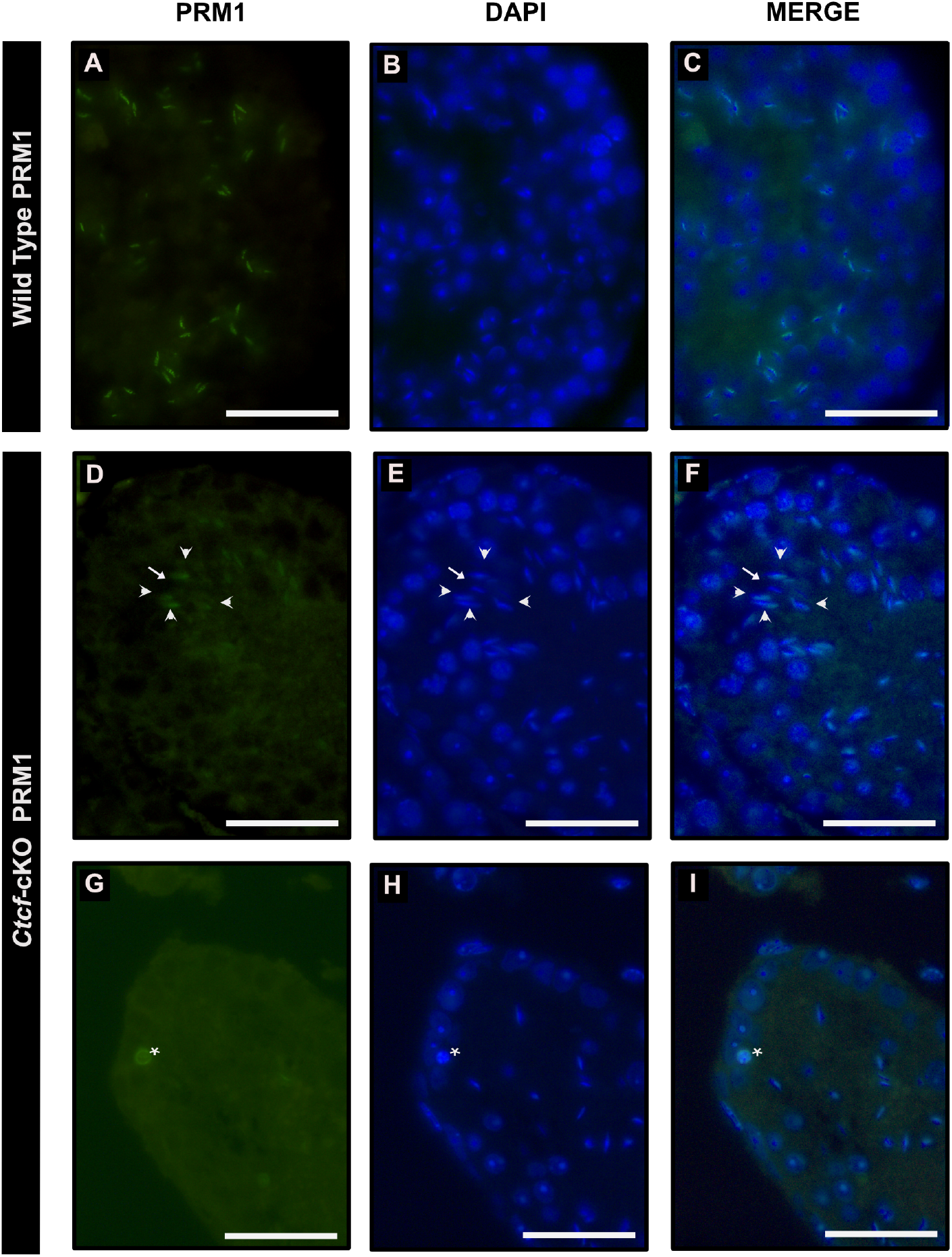
Immunofluorescent staining pattern of PRM1 in Wild Type and *Ctcf*-cKO testis sections. (**A-C**) Show immunodetection of PRM1 in elongated spermatids from WT testis sections. Some representative elongated spermatids with the signal of PRM1are pointed out with arrows. (**D-F**) Immunodetection of PRM1 in elongated spermatids from *Ctcf*-cKO testis sections. Elongated spermatids with the signal of PRM1are pointed out with arrows. (**G-I**) Immunodetection of PRM1 in elongated spermatids from *Ctcf*-cKO testis sections. 30% of the analyzed seminiferous tubules contained cells with PRM1 expression and abnormal morphology.

## Discussion

Although CTCF is a key molecule for sperm formation, fertility, and sperm epigenome establishment in mice (Jung et al., 2017), little is known regarding its function throughout the different stages of spermatogenesis. In this study, by analyzing the transcriptome of single cells at different stages of spermatogenesis, we found that there are no major alterations in the biological functions of mitotic and meiotic cells under the absence of CTCF. However, haploid cells in spermiogenesis undergo significant changes in their transcriptional profiles and morphology development. We found that early-round spermatids are the most affected cell type, yet they still maintain certain identity that allowed us to map them in our scRNA-seq analysis. Our GSEA enrichment analysis of the mis-regulated genes indicated that early-round spermatids have problems related to spermatids development and sperm morphology. When then looked into the spermatids morphology and found defects in acrosome biogenesis throughout the different stages of spermiogenesis (i.e., round, elongating and elongated spermatid stages). Furthermore, we also found elongating and elongated spermatids with defects in nuclear compaction and protamine incorporation. In a previous work, we reported mature sperm from the cauda epididymis from *Ctcf*-cKO mice with acrosome and nuclear compaction malformations (Hernandez-Hernandez et al., 2016).

In this work, by scRNA-seq we underscored that these defects are produced by alterations in transcriptional profiles of related genes in early-round spermatids. Single-cell RNAseq studies in mouse spermatogenesis have shed light on the heterogeneity found during this process (Green et al., 2018; Hermann et al., 2018). Thus, we used the stage-specific markers elucidated by these studies to identify and validate cluster phenotypes within our bioinformatics pipeline. Furthermore, it is well documented that scRNA-seq can underscore cell phenotypes that are otherwise masked when RNA-seq bulk analysis from whole tissues is performed (Bacher and Kendziorski, 2016; Chaudhry et al., 2019). Indeed, we found that during the early round stage, promoter DNA methylation is also altered in the absence of CTCF. DNA methylation in the germline has the potential to regulate gene expression in the next generation. Hence, failures in this process lead to serious consequences for post-fertilization development (Stewart et al., 2016). In the mouse, genome-wide demethylation occurs early in the development of primordial germ cells. However, a wave of de novo methylation imprints epigenetic memory at specific genomic sites (imprinted regions) at the prospermatogonia stage before birth (McSwiggin and O’Doherty, 2018; Reik et al., 2001).

We found the promoter DNA methylation pathway to be affected in round spermatids of the *Ctcf*-cKO, implying that in a WT scenario this biological pathway is an ongoing process. Yet, de *novo* DNA methylation during round spermatid development has not been reported. Instead, a mechanisms of active DNA methylation has been suggested to be acting on order to repress transposon in the male germline (MORC1 represses transposable elements in the mouse male germline (Pastor et al., 2014). Accordingly, the gene responsible for this transposon repression, MORC1, is also downregulated in mid-round spermatids from *Ctcf-*cKO mice (Table S6). Therefore, the biological implications for alterations in DNA methylation and transposon repression at the haploid stage, remain to be elucidated.

Although in low numbers, sperm with several morphological and biochemical alterations are still produced in the absence of CTCF (Hernandez-Hernandez et al., 2016), most of these defects are created at the transcriptional level in early round spermatids, and the resulting phenotype is observed throughout the whole process of sperm differentiation (this work). However, the exact mechanism by which CTCF is controlling these biological functions has not been fully addressed. A possible mechanism may be related to the establishment of the sperm epigenome at the same time that histones are replaced in most of the genome and retain at specific genomic regions at the stage of round spermatids (Jung et al., 2017). At this stage, CTCF and cohesin complexes occupy specific genomic regions orchestrating the organization of the genome or sperm epigenome (Jung et al., 2019; Jung et al., 2017; Torres-Flores and Hernandez-Hernandez, 2020). Thus, we speculate that absence of CTCF in round spermatids may have a strong impact on sperm epigenome organization by altering sperm-specific transcriptional regulatory programs. Detailed chromatin accessibility studies at the single cells level are needed to explore this hypothesis. Our work highlights the importance of CTCF for the proper progression of spermiogenesis.

## Funding

Universidad Nacional Autónoma de México-PAPIIT grant IN225917 Hospital Infantil de México Federico Gómez grant HIM/2018/079 SSA 1518 Instituto Nacional de Medicina Genómica project ID348 Direct funding of Patronato del Hospital Infantil de México Federico Gómez.

## Author Contributions

UTF and AHH designed the study. UTF, TLS and AHH performed sample preparation. UTF and FDE performed scRNA-seq libraries prep and sequencing data processing. FDE and RRV contributed to the NGS process. ROH and AHH performed tissue preparation for immunofluorescence and electron microscopy.

AVJ and ORA performed sequencing data pre-processing, and enrichment analysis. IJT and UTF performed the rest of the data analysis. IARV input with scientific ideas. OME, GHVN, IARV, ORA, MCGR and AHH contributed with reagents, equipment and funding. AHH and ORA supervised the experiments and data analyses. AHH and UTF wrote the manuscript. All the authors discussed the data and reviewed the manuscript.

## Competing Interests

The authors declare that they have no conflict of interest.

## STAR methods

### Animals

We have previously generated and validated a transgenic mouse strain that undergoes depletion of CTCF during spermatogenesis (Hernandez-Hernandez et al., 2016). For this work, we used male mice of 12 to 15 weeks old to minimize age related variations. We compared *Ctcf* conditional knockout mice (*Ctcf*-cKO, with a Stra8-iCre-*Ctcf* ^f/Δ^ genotype) versus Wild type (WT) littermates with genotypes *Ctcf*^f/f^, *Ctcf*^wt/f^ or *Ctcf*^wt/wt^.

### Single cell RNA sequencing

We performed spermatogenic cell dissociation from testes as previously reported (Kossack et al., 2013) with some modifications. Briefly, we extracted and removed the tunica albuginea from the testes of two WT and two *Ctcf*-cKO mice. We gently spread seminiferous tubules on a small petri dish and transferred them into a 15 mL conic tube containing 5 mL of enzymatic buffer (20 mM Hepes pH 7.2, 6.6 mM sodium pyruvate, 0.05% lactate in 1× HBSS), supplemented with 100 U/mL collagenase type I and 2 U/mL of DNase. Tubules were incubated for 25 minutes at 32°C and the suspension was filtrated throughout a 40 μm nylon mesh. FBS was added to the filtrated solution to a final concentration of 2% collecting. The filtrates of two tubes from the *Ctcf*-cKO mouse were pooled in a new tube. Then, we pelleted the cells by centrifugation at 1000 g for 10 min at 4°C, we carefully aspirated the supernatant and added 1 mL of HBSS enzymatic digestion buffer. We then mixed thoroughly and transferred the suspension into a 1.5 mL tube. We pelleted cells by centrifugation at 1000 g for 10 min at 4°C and we aspirated the supernatant. To eliminate as much as possible the erythrocytes from the cell suspension, we washed the pellet three times with ice cold PBS. Finally, we pelleted the cells by centrifugation at 1000 g for 10 min at 4°C and we removed the supernatant. The spermatogenic cells were re-suspended in PBS plus 0.2% BSA and cell concentration was adjusted to 2.5 × 10^6^ cells/mL. Only cell suspensions with a viability greater than 90% were used for downstream analysis.

We performed single cell isolation and cell barcoding with the Droplet-based Single-cell isolator (one-touch ddSEQ, Bio-Rad) according to the manufacturer’s instructions. We subsequently prepared Single-cell-barcoded RNA-Seq libraries using Nextera technology included in the SureCell WTA 3′ Library Prep Kit (Illumina). Next generation sequencing was performed in a NextSeq (Illumina) with an attainable depth of 250,000 reads/cell. Four data sets from two different WT and two different *Ctcf*-cKO mice were generated.

### Data analysis

In order to process single cell mRNA sequencing data, we generated the count expression matrixes by aligning the raw data to the mouse genome (GRCm38.98) following previous reports (Romagnoli et al., 2018; Tian et al., 2018), using the default parameters. For downstream analysis of the count expression matrixes, we used the SeqGeq v1.6.0 software (BD Life Sciences, Ashland, OR, USA.). Within SeqGeq, we performed quality control to identify non-outlier parameters meeting a threshold of 8 units dispersion or greater, we also identified that all cells displayed less than 8% of mitochondrial RNA. Quality control of cells was confirmed prior downstream data analysis. Next, we generated graph-based clustering using the Seurat algorithm (Satija et al., 2015), with the top 2070 most highly dispersed genes and non-outlier cells followed by dimensionality reduction through PCA guided UMAP (Becht et al., 2018). We then identified clusters using their specific gene expression as previously reported in other scRNA-seq datasets (Green et al., 2018; Hermann et al., 2018). For differential expression analysis we used a +/− 2 fold-change and a FDR adjusted p-value < 0.05. Finally, we used the trajectory inference algorithm Monocle 2 (Trapnell et al., 2014) for deeper exploration of the specific clusters in either genetic condition (WT vs *Ctcf*-cKO).

All the single-cell RNA-seq data has been deposited in the GEO portal and the accession number will be release upon acceptance of this paper, however a private access token may be provided under request

### Pathway Enrichment Analysis

To analyze functional annotation of the up- and down-regulated genes in our dataset we used DAVID v6.8 (The Database for Annotation, Visualization and Integrated Discovery) (Huang da et al., 2009). For pathway enrichment analysis we used the GSEA computational method (Gene Set Enrichment Analysis), (Subramanian et al., 2005) available at http://software.broadinstitute.org/gsea/. We obtain and used all the gene set annotations from the GSKB database available at: http://ge-lab.org/gskb (Lai,2016). Statistical significance was assigned with an FDR ≤ 0.05 and p-value ≤ 0.01.

### Enrichment Maps

We constructed enrichment maps with the pathway enrichment analysis information in Cytoscape software. Using a Jaccard index values ≥ 0.4 we associated statistically significant enriched pathways according to their shared genes. We set the Biological function collections tags with the AutoAnnote app.

### Histological Sections processing

We performed electron microscopy and immunofluorescent analysis as previously reported (Hernandez-Hernandez et al., 2016).

## Notes

### Competing Interest Statement

The authors have declared no competing interest.

