## Supplementary Figures for "Spermiogenesis alterations in the absence of CTCF revealed by single cell RNA sequencing"

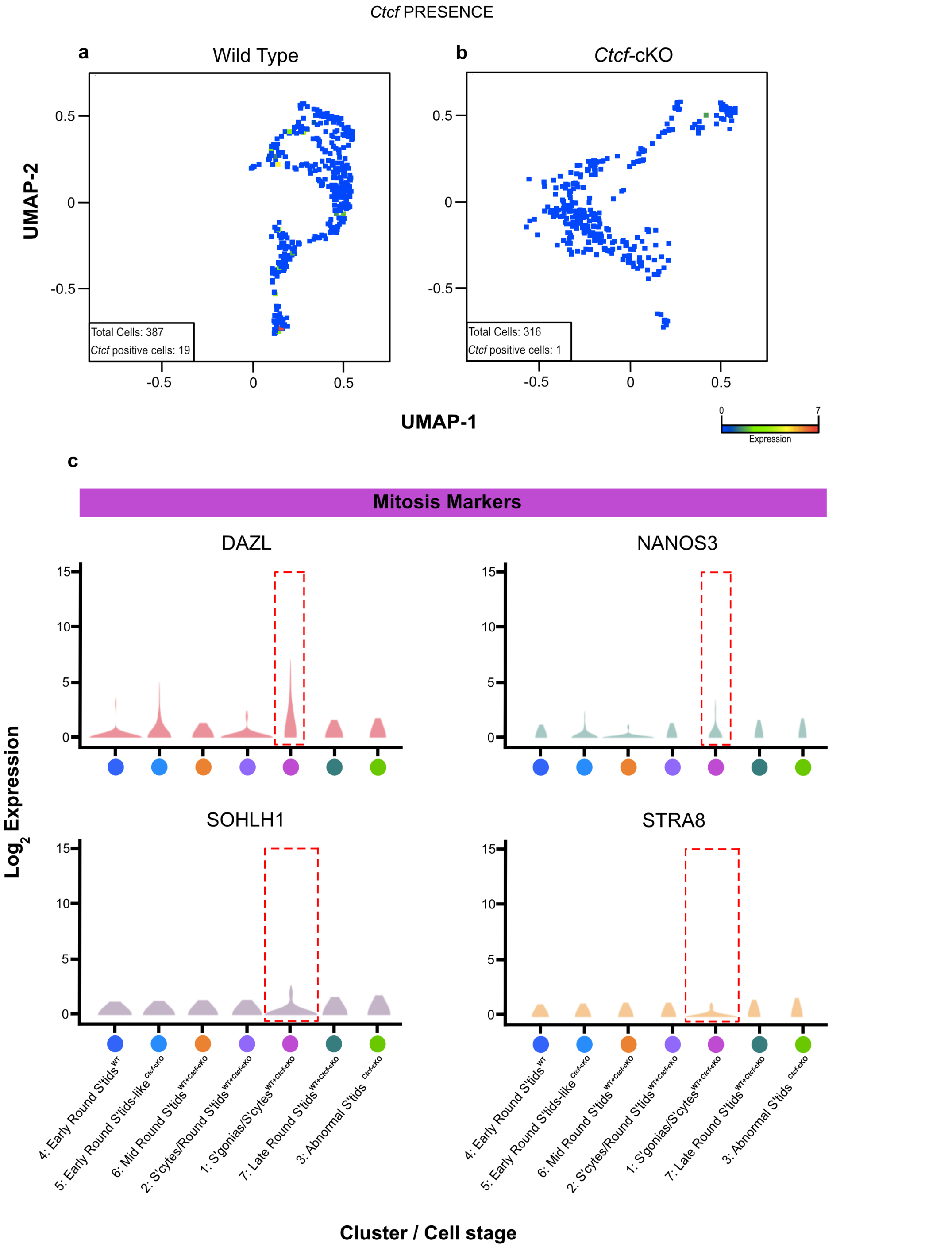


**Supplementary Figure 1. Distribution of key genes.** (**A** and **B**) show UMAP projection displaying the expression of *Ctcf* in WT and *Ctcf*-cKO spermatogenesis cell clusters. Number of cells with expression of *Ctcf* are indicated within each panel. (**C**) Violin plots displaying the distribution of the expression levels of some mitotic genes. Although cluster 1(S'gonias/S'cytes) (dotted rectangle) show greater expression levels of the four markers, no significative differences were identified (Kruskal-Wallis test, $p>0.05$).

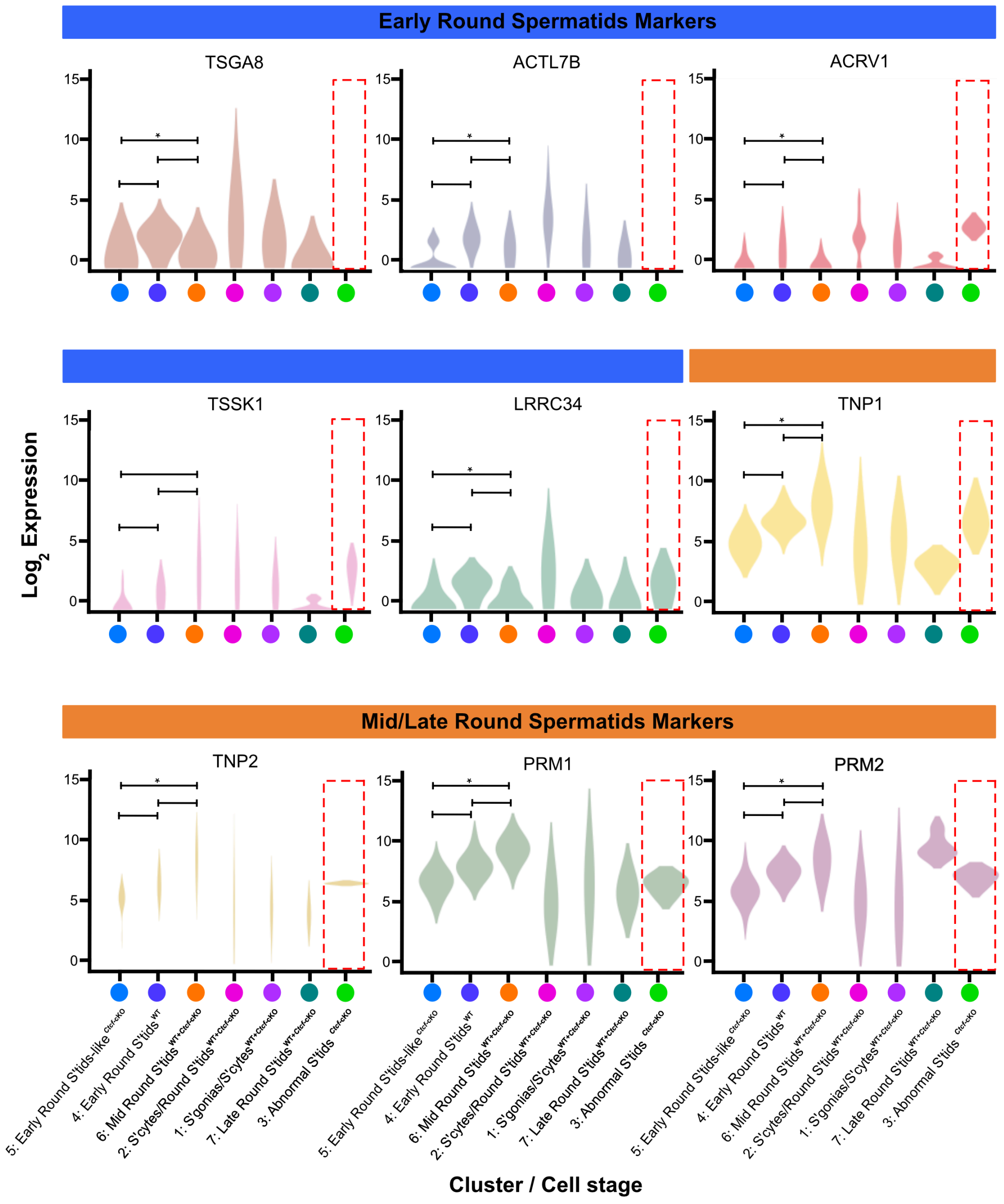


**Supplementary Figure 2. Violin plots showing the distribution of the expression levels of spermiogenesis genes across all the cell clusters from WT and *Ctcf*-cKO mice**. Dotted red line rectangle highlights cluster 3 (abnormal spermatids specific from *Ctcf*-cKO mice) displaying abnormal expression of all the analyzed genes. Distribution of the expression levels of all the markers in cluster 5 (specific from the *Ctcf*-cKO mice) are similar to cluster 4 (Early-round spermatids, specific for WT) but not to cluster 6 (mid-round spermatids) (Kruskal-Wallis, $p>0.05$).

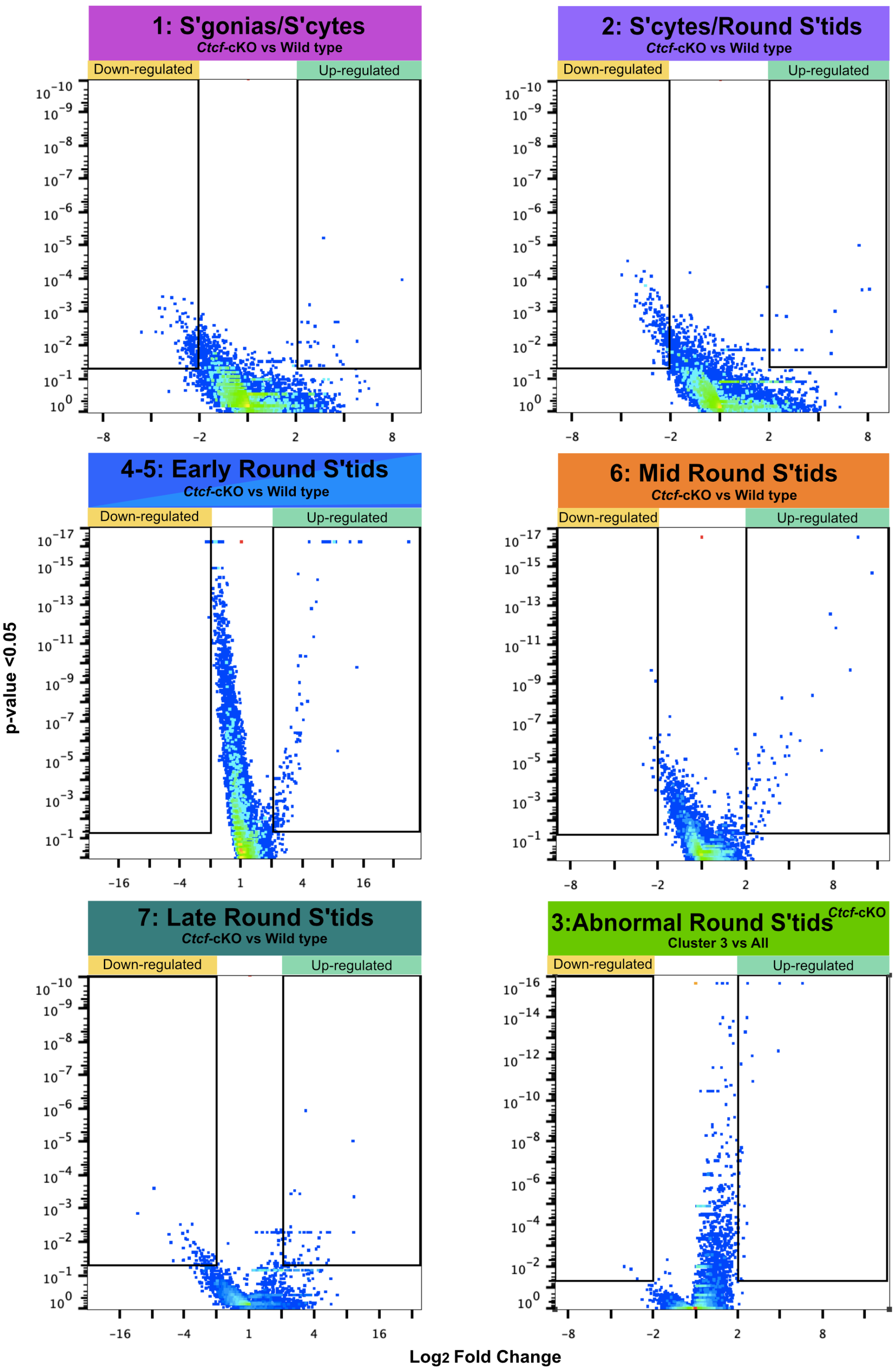


**Supplementary Figure 3. Volcano Plots**. Each panel display the down- and up-regulated genes in *Ctcf*-cKO cell clusters with a +/- 2-fold-change and FDR adjusted p-value < 0.05. Comparisons between *Ctcf*-cKO versus WT were done using cells within the same cluster (clusters 1, 2, 6 and 7), whereas cluster 5 (from *Ctcf*-cKO) was compared to cluster 4 (from WT) and cluster 3 (from *Ctcf*-cKO) was compared to all the clusters.

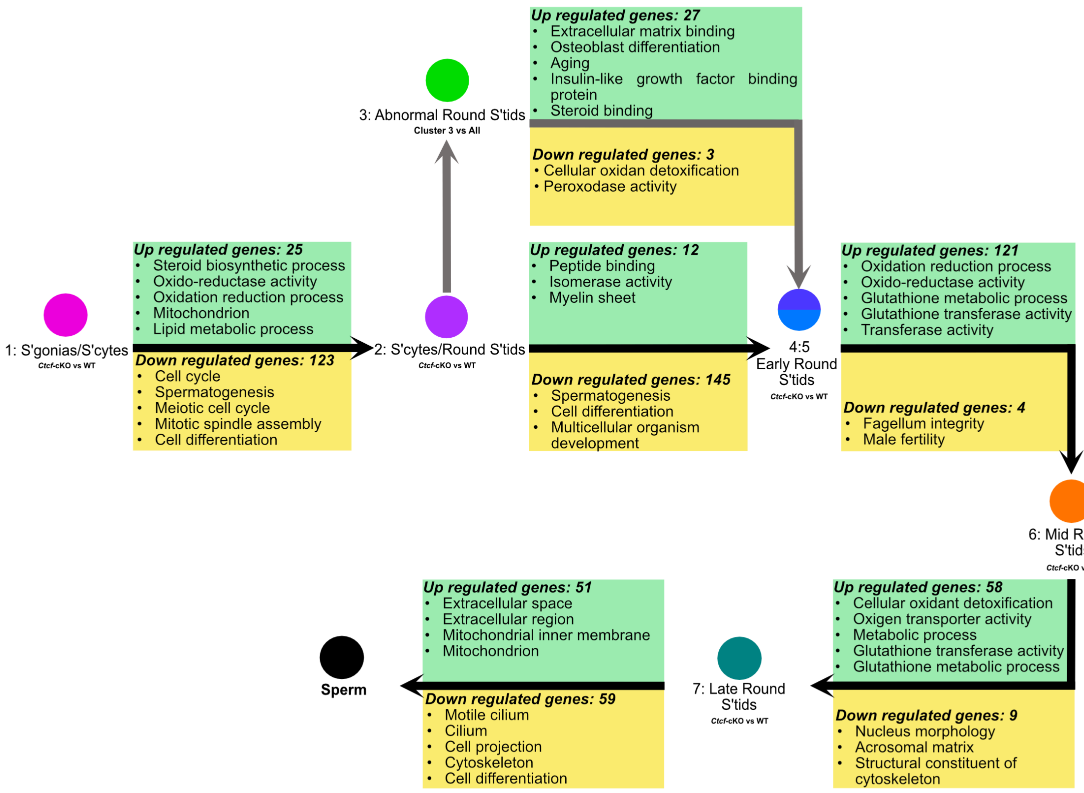


**Supplementary Figure 4. Functional annotations from the differentially expressed genes in *Ctcf*-cKO condition.** Comparisons between *Ctcf*-cKO versus WT were done using cells within the same cluster (clusters 1, 2, 6 and 7), whereas cluster 5 (from *Ctcf*-cKO) was compared to cluster 4 (from WT) and cluster 3 (from *Ctcf*-cKO) was compared to all the clusters. The diagram shows the first five GO terms identified with DAVID v6.8. For differential expression analysis we used a +/- 2-fold-change and a FDR adjusted p-value < 0.05. Green rectangles highlight the up-regulated genes while yellow rectangles the down-regulated genes.
